# A mobile CRISPRi collection enables genetic interaction studies for the essential genes of *Escherichia coli*

**DOI:** 10.1101/2023.04.21.537703

**Authors:** Kenneth Rachwalski, Megan M Tu, Sean J Madden, Drew M Hansen, Eric D Brown

## Abstract

Ordered collections of precise gene deletions in *E. coli* and other model microbes have proven to be invaluable resources to study the dispensability of the non-essential genome in different environmental and genetic contexts. The dispensability of canonically essential genes, however, has been largely unstudied in such contexts. Advances in gene editing, in particular CRISPR interference (CRISPRi), have enabled depletion of essential cellular machinery to study the downstream effects on bacterial physiology. Here, we describe the construction of an ordered *E. coli* CRISPRi collection, designed to knock down the expression of 356 essential genes with the induction of a catalytically inactive Cas9, harbored on the conjugative plasmid pFD152, that also encodes for expression of specific guide RNA scaffolds. This mobile CRISPRi library can be conjugated into other ordered genetic libraries, to assess combined effects of essential gene knockdowns with non-essential gene deletions. As proof of concept, we probed cell envelope synthesis with two crosses. First, we crossed a deletion in the gene *lpp* with the entire CRISPRi library. Similarly, we crossed the CRISPRi knockdown of *lolA* with ∼4,000 deletion strains of the Keio collection. These experiments revealed a number of notable genetic interactions for the essential phenotype probed and, in particular, showed supressing interactions for the loci in question.

## Introduction

Essential genes are defined as those that are required for the survival of an organism under normal growth conditions. The environmental and genetic context of a bacterium will establish which biological processes are required for growth and survival^1,2^; gene essentiality is not a binary determination. In *Escherichia coli*, approximately 300-400 genes have been identified as being essential for growth in standard laboratory media using a variety of approaches^3–6^. These genes perform vital functions required for bacterial survival and are thereby sought-after targets for antibacterial drug discovery efforts^7^. Indeed, patterns and paradoxes in gene dispensability offer profound insight into gene function and can be leveraged for small-molecule screens that search for novel anti-bacterial compounds^8–10^.

An important tool to study the dispensable gene set in *E. coli* is the Keio collection of ∼4,000 gene deletion strains^4^. The Keio collection has been extensively screened to identify the environmental conditions, for example, different growth media and chemical stressors, where canonically non-essential genes are rendered essential for *E. coli* growth^11–13^. Likewise, millions of unique double deletion strains have been generated with the Keio collection and screened using synthetic genetic array technology to uncover genetic contexts where genes become unexpectedly essential^14–22^. In contrast, there is great difficulty in performing such large scale gene-gene interaction assays that include essential processes in *E. coli*. Chemical and antibiotic probes are often used to investigate the effects of perturbing essential processes in different genetic backgrounds, in what are described as ‘chemical genomics’ screens^11,12,23,24^. Unfortunately, existing chemical probes target a small subset of essential biology, or, work only in bacterial strains with permeabilized membranes or deficient efflux systems. As such, similar to synthetic genetic arrays for dispensable genes, a genetics-based approach to disrupt essential genes in existing mutant collections would be highly advantageous to the study of genetic interactions with the essential genome.

Recently, focus has shifted to the use of CRISPR interference (CRISPRi) to study the effects of perturbing essential genes and processes^25,26^. CRISPRi enables inducible and titratable knockdown of gene expression using a catalytically inactive Cas9 (dCas9) which is guided to a target gene by a 20-nt single guide RNA sequence (sgRNA), thereby blocking the transcription of nascent mRNA^25^. The versatility of CRISPRi has enabled investigations of gene essentiality in *E. coli*^5^ and other non-model microbes^27^. The depth of *E. coli* annotation as a model microbe^28^ makes it ideal for studying biological nuances using genetic tools. In *E. coli*, genome-wide CRISPRi fitness screens are commonly conducted using pooled collections of sgRNAs ^5,29–32^. In these screens, gene knockdown by CRISPRi is induced in the pooled collection, which is then grown for a pre-determined number of generations before the relative abundance of different sgRNAs is quantified by next generation sequencing. Pooled approaches have also been used to conduct gene-gene interaction studies with *E. coli* deletion strains^33^. However, this approach requires the *de novo* construction of a gene deletion in a genetic background expressing a chromosomal dCas9 and is thereby not immediately compatible with existing arrayed genomic collections such as the Keio collection^4^. Alternatively, ordered collections can be screened for individual strain fitness without worrying about cross-feeding and other phenomena often observed with pooled mutant screening approaches^24,34^. Screening ordered collections also allows for investigating the effects of knocking down an individual gene on bacterial cell physiology– for instance changes in cell morphology^35,36^, the cellular metabolome^37,38^, or the cellular proteome^38,39^. While an ordered *E. coli* CRISPRi collection has been published recently by Silvis *et al* ^35^, the tri-parental mating described to introduce *dcas9* and the sgRNAs into new strains was reported to have a cross-contamination rate of ∼30%, making this approach unsuitable for creating large numbers of mutants, required for performing genome-wide genetic interaction studies.

Here, we describe an ordered *E. coli* CRISPRi collection of 356 sgRNAs targeting essential genes built on the conjugative plasmid pFD152^40^. Unlike existing ordered CRISPRi libraries, we were able to rapidly and efficiently move our collection into any genetic background of interest by high-throughput conjugation, allowing for the interrogation of genetic interactions between the dispensable and indispensable genome of *E. coli* K-12. As a proof of principle, we conjugated our CRISPRi collection into a strain deleted for gene *lpp*, encoding Braun’s lipoprotein. Lpp is the most prevalent lipoprotein in the cell, and has well characterized synthetic viable interactions with the essential genes involved in lipoprotein transport^41,42^. As expected, we found that CRISPRi constructs targeting essential genes involved in lipoprotein processing and transport (*lnt, lolABC*) showed considerably less growth inhibition in *E. coli* Δ*lpp*. In line with this observation, we also demonstrated that conjugating pFD152:*lolA* into the Keio collection of ∼4,000 deletion strains identified Δ*lpp* as one of the strongest suppressors of growth inhibition resulting from knocking down *lolA* expression. The platform has the potential to enhance our understanding of the biology underpinning essential processes in *E. coli*, and can provide biological insights to design unconventional screening platforms aimed at identifying novel chemical inhibitors of essential processes.

## Results & Discussion

To facilitate genetic interaction studies for the essential genes of *E. coli*, we sought to create a collection of essential gene knockdowns constructed on a mobile CRISPRi plasmid using the recently published pFD152 vector^40^. CRISPRi knockdown by pFD152 is inducible with anhydrotetracycline (aTc), which tightly controls the expression of *dcas9* allowing for titratable silencing of gene expression^40^. Additionally, pFD152 contains a constitutively expressed sgRNA, an RP4 origin of transfer that enables rapid conjugation into target strains, an origin of replication compatible with most of the commonly used plasmids in *E. coli*, and a spectinomycin selectable marker that is distinct from those commonly used in existing *E. coli* ordered deletion collections **(Figure 1A**). We used the CRISPRbact tool to design sgRNAs that minimize off-target binding commonly seen with sgRNAs in CRISPRi assays^30,31^. We then introduced these sgRNAs into pFD152 using Golden-Gate assembly^40^ and confirmed all plasmid sequences by Sanger sequencing. A comprehensive list of all 356 sgRNAs in our collection can be found in **Supplementary Table 1**. Of the 356 genes represented in our collection, 353 were selected based on having an essential classification in a recent transposon sequencing survey of gene essentiality^3^ and three non-essential genes were included as controls (*waaU, lpxL*, and *bioA)*. Strains harboring sgRNAs targeting specific genes were arrayed at 384-density with 17 randomly dispersed strains carrying the pFD152 empty vector as a control, which has a 20 nt spacer sequence *in lieu* of a gene-specific sgRNA (**Figure 1A)**.

**Figure 1.**
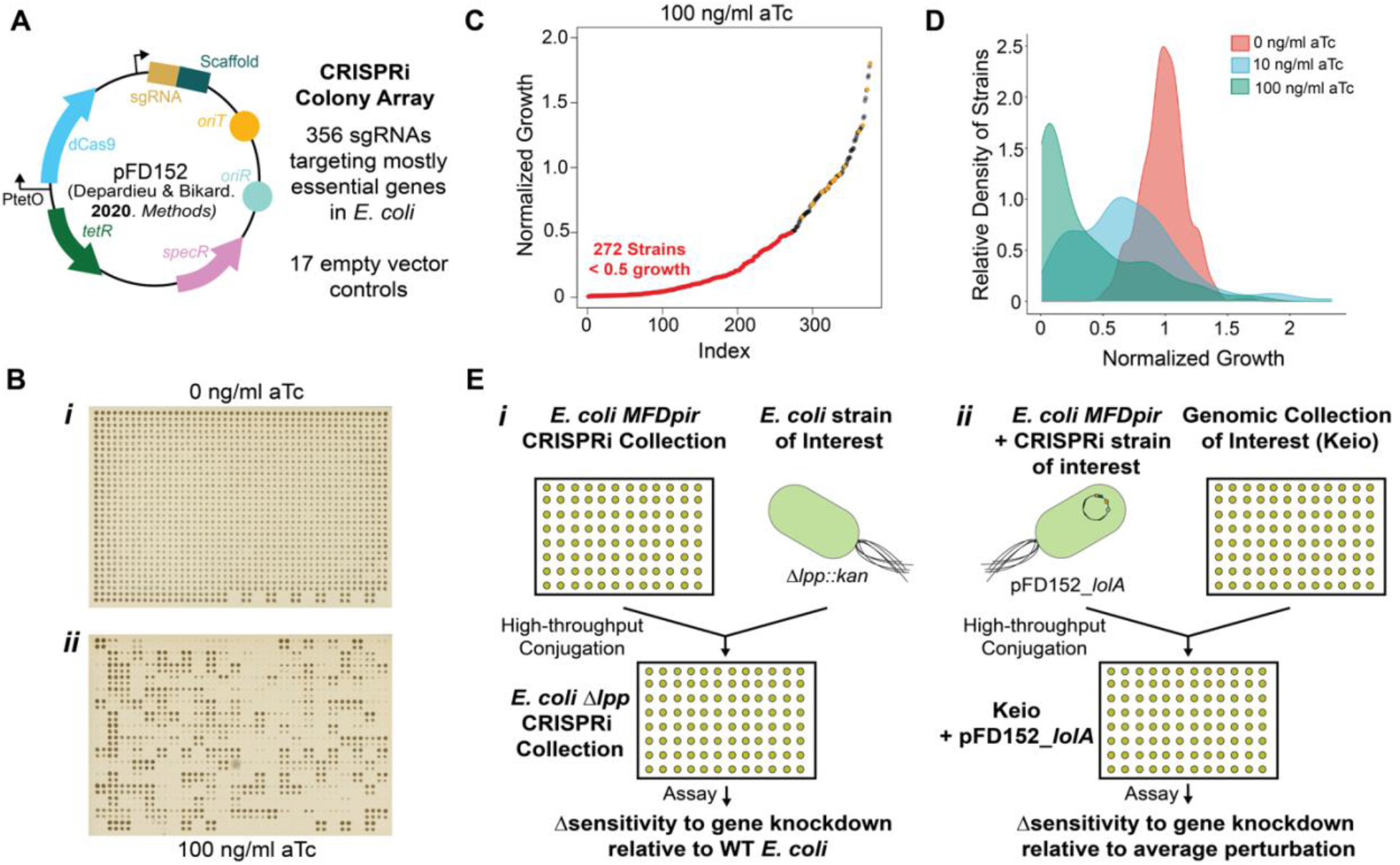
Overview of the *E*. coli mobile arrayed CRISPRi Collection. **(A)** The previously published conjugative CRISPRi plasmid pFD152^40^, contains all required components for anhydrotetracycline(aTc) inducible gene silencing in *E. coli*. 356 sgRNAs targeting essential genes were cloned into pFD152 and validated by Sanger sequencing, and transformed into WT *E. coli BW25113* and the conjugative donor strain *E. coli* MFD*pir*. Transformants were arrayed along with 17 strains carrying an empty vector that served as controls. **(B)** Shown are images of the CRISPRi collection grown at high-density in a colony array in the absence (*i*) or in the presence of (*ii*) aTc. Shown is an array grown at 1,536 colony density wherein each strain was grown in technical quadruplicate. **(C)** A rank-ordered plot of mutant growth at 100 ng/ml aTc. 272 strains in the CRISPRi array demonstrated a substantial growth defect (normalized growth <0.5) upon CRISPRi induction. Empty vector controls are represented by points labeled in orange. **(C)** A density plot representing the distribution of growth of the CRISPRi collection at different aTc concentrations. Collection-wide growth inhibition was dose dependant on concentration of inducer. **(D)** Schematic representing two complementary uses of the mobile arrayed CRISPRi collection to conduct gene-gene interaction studies with essential genes. *(i)*The entire mobile CRISPRi array could be rapidly conjugated into a deletion strain of interest using high-throughput conjugation to investigate the effects of a gene deletion on the sensitivity to disrupting different essential processes, or, *(ii*) an individual CRISPRi plasmid could be introduced into existing genomic collection such as the Keio collection, to identify genetic backgrounds where disrupting a specific essential process is enhanced or suppressed.

When grown in the presence of aTc which induces CRISPRi knockdown, a growth defect was observable for a large portion of our collection (**Figure 1B**). With 100 ng/ml aTc added to the growth media, 272 from the 356 strains showed at least a 50% reduction in growth (**Figure 1C**). We screened the CRISPRi collection for growth inhibition in both rich (LB) and minimal (MOPS minimal) microbiological media at varying aTc concentrations and observed a dose-dependent growth defect (**Figure 1D, Figure S1, Supplemental Table 2**) while maintaining consistency between screening replicates (**Figure S2**). We note that several of our CRISPRi constructs did not cause a growth defect despite targeting the expression of an essential gene, however, these genes largely overlap with previously reported sets of essential genes which do not elicit a growth defect when perturbed by CRISPRi^5^. We speculate that the products of these genes may have long half-lives or that the bacterium may tolerate reduced levels of viable protein. Nonetheless, there is merit to including these strains in our collection to identify genetic or environmental contexts where a growth defect might be elicited.

For ease of moving the collection into different genetic backgrounds, we transformed each CRISPRi knockdown plasmid into the conjugative donor strain *E. coli* MFD*pir*^43^, creating an ordered mobile CRISPRi collection. Here, we present two different applications of the mobile CRISPRi collection (**Figure 1E**). First, the complete CRISPRi collection can be conjugated into a specific genetic background of interest, for example, a deletion mutant of the Keio collection. By comparing the growth inhibitory effects of each CRISPRi construct in the mutant background to that of the wildtype background, we can uncover CRISPRi knockdowns of essential processes that exhibit enhanced or suppressed growth inhibition in the mutant background. Second, we can conjugate an individual CRISPRi knockdown strain into existing *E. coli* genomic collections, for instance into the Keio collection. This allows testing genetic interactions between an essential gene and all the ∼4,000 genes in the Keio collection simultaneously, supplementing existing ‘synthetic genetic array’ screening datasets^14,17^. Further, due to the compatibility of pFD152 with commonly used *E. coli* plasmids, individual CRISPRi constructs could also be conjugated into plasmid-borne arrayed genomic collections including transcriptional reporter collections^44^ and protein over-expression collections^45^.

As a proof of principle, we elected to probe cell envelope synthesis in *E. coli*. First, we conjugated the mobile CRISPRi collection into *E. coli* Δ*lpp*. Lpp is the most abundant lipoprotein in *E. coli* and acts to covalently tether peptidoglycan to the outer membrane^46–49^. Perturbing essential genes involved in lipoprotein transport, namely the Lol system^50^, results in the mis-localization of lipoproteins, thereby causing cell death^50–52^. In the absence of Lpp, the growth defect associated with disrupting lipoprotein transport is reduced^42^. As such, we expected to observe suppressed killing by CRISPRi knockdowns targeting lipoprotein synthesis and trafficking upon mating the CRISPRi plasmids into an *lpp* deletion strain. Following previously published high-throughput conjugation workflows^21^, we introduced the CRISPRi collection into *E. coli* Δ*lpp* (**Figure S3**). The wildtype (*lpp+*) and Δ*lpp* CRISPRi collections were then screened at varying aTc concentrations on minimal media, and the growth observed for each strain in the Δ*lpp* background was normalized to that of the wildtype (*lpp+*) collection at each concentration of inducer (**Figure S4**). In the absence of inducer both the wildtype (*lpp+*) and Δ*lpp* collection grew comparably, however, with induction we observed CRISPRi constructs that exhibited either enhanced or suppressed growth inhibition in the Δ*lpp* background (**Figure 2A, Supplemental Table 3**). Using an arbitrary threshold of 5-fold change in growth between the Δ*lpp* and wildtype (*lpp+*) collections, growth inhibition by 17 CRISPRi constructs was enhanced and for 42 CRISPRi constructs was suppressed in at least one concentration of inducer. Interestingly, more than half (24/42) of the suppressors were identified at the lowest concentration of aTc (5 ng/ml), but growth inhibition with all of these low induction suppressors was restored at higher concentrations of inducer (aTc > 10 ng/ml). Looking at 100 ng/ml aTc specifically, it’s unsurprising that the four strongest suppressors identified were CRISPRi constructs targeting essential genes involved in lipoprotein synthesis and trafficking – *lolA, lolB, lol*C, and *lnt* (**Figure 2B**). The 17 CRISPRi plasmids that exhibited enhanced growth inhibition in an Lpp deficient background were mostly identified at higher concentrations of aTc (>10 ng/ml). These enhancers are involved in the biosynthesis of essential phospholipids (*gpsA, psd, plsBC, pssA*), the outer membrane (*bamA, lpxC, lptA, kdsC*), and in cell division (*ftsAHWY, dicA, zipA*). Fortunately, some of these interactions were also testable with available chemical inhibitors of these essential processes (**Figure 2C**). Using a commercially available inhibitor of the Lol system^53^ (LolCDE-in-1) and of BamA^54^ (MRL-494), we showed that a Δ*lpp* strain was less susceptible to the inhibitor of the Lol system, and more susceptible to the BamA inhibitor as predicted in our screen.

**Figure 2.**
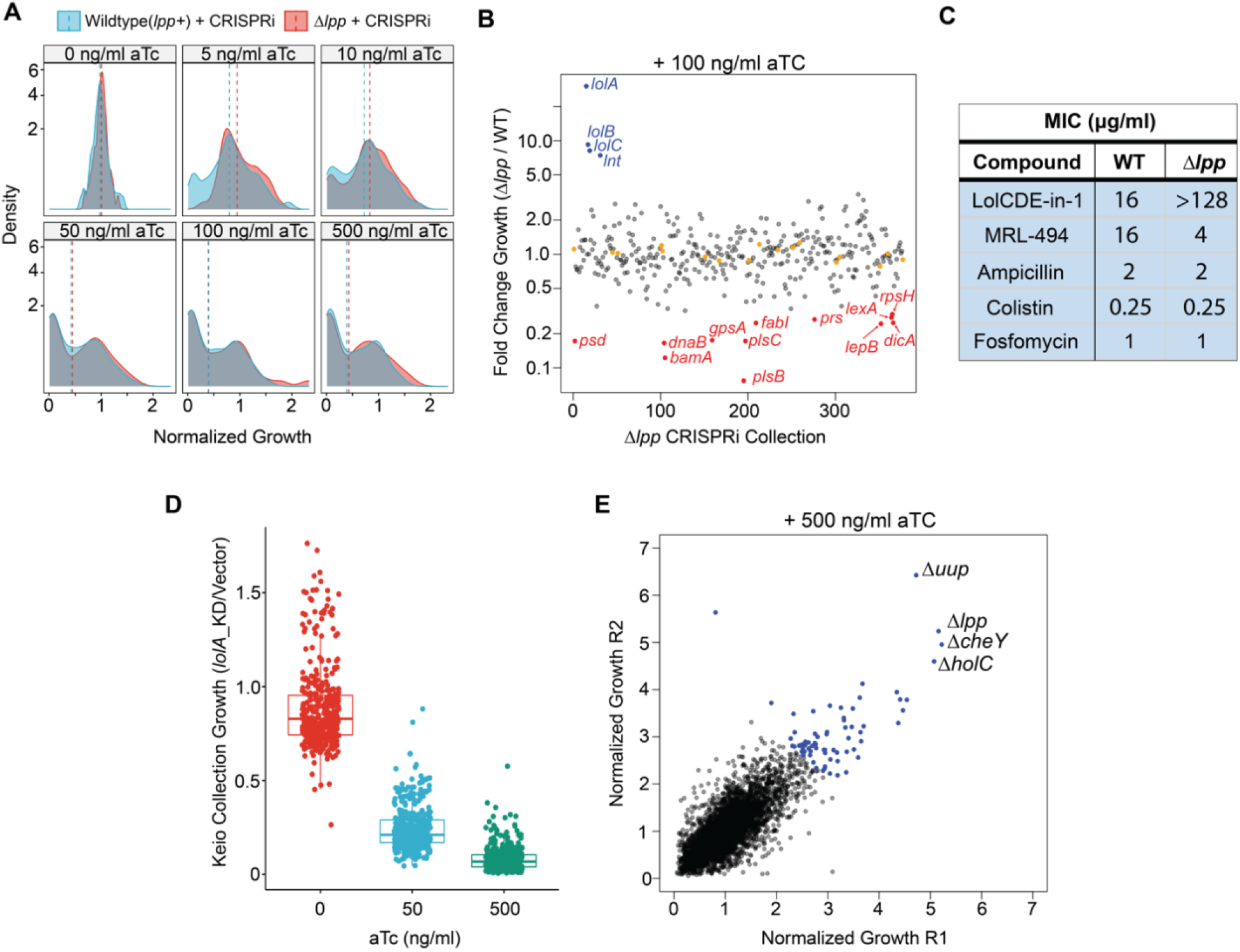
CRISPRi genetic interaction screens identify a known genetic interaction between Lpp and lipoprotein transport. **(A)** The mobile arrayed CRISPRi collection was introduced into *E. coli* Δ*lpp*, and the collection was screened at 6 different concentrations of aTc in minimal media. Density plots show the distribution of growth for both the WT and Δ*lpp* collections at different levels of CRISPRi induction. Average growth of the WT and Δ*lpp* collections is represented by the dotted lines in each density plot. **(B)** The fold-change in growth of the Δ*lpp* CRISPRi collection relative to the wildtype (*lpp*+) CRISPRi collection shown at 100 ng/ml aTc. CRISPRi constructs with suppressed growth inhibition and shown in blue, and CRISPRi constructs with enhanced growth inhibition are shown in red. Empty vector controls are shown in orange. Hits were determined as data points that deviate by at least three-standard deviations from the mean. **(C)** Minimum inhibitory concentrations (MICs) of different compounds against Wildtype and Δ*lpp E. coli* in minimal growth media. **(D)** The *lolA* knockdown plasmid was introduced into all ∼4,000 mutants in the Keio collection by high-throughput conjugation, and the Keio collection was screened at 0, 50, and 500 ng/ml of aTc. Shown is the change in growth between mutants harboring the *lolA* knockdown plasmid compared to those same mutants harboring the empty vector from a representative plate of the Keio collection (384 strains). **(E)** A replicate plot showing the normalized growth of two biological replicates of the pFD152*_lolA* Keio collection at 500 ng/ml aTc. Mutants which demonstrated growth inhibition suppressed by at least 3 standard deviations from the mean of the dataset are coloured in blue.

By screening at different inducer concentrations we could discriminate between different degrees of suppression and enhancement. For instance, the suppressive interaction of Δ*lpp* on growth inhibition by pFD152:*lolA* was observed at concentrations greater than 50 ng/ml of aTc, whereas suppression by pFD152:*lspA* was only observed at the two lowest concentrations of inducer – 5 and 10 ng/ml aTc (**Figure S5**). Both *lspA* and *lolA* are involved in lipoprotein synthesis and trafficking respectively, and Lpp deficient bacteria are known to be resistant to depletion or inhibition of these proteins^42,55,56^. Likewise, enhanced growth inhibition by pFD152:*zipA* was observed at 10 ng/ml aTc whilst that by pFD152:*plsC* occurred at concentrations greater than 50 ng/ml aTc. We further confirmed the results from our Δ*lpp* CRISPRi screen using a standard dilution plating assay (**Figure S6**). These data suggested the need to screen CRISPRi collections at multiple concentrations of inducer to identify all possible gene-gene interactions. Identifying *bona-fide* synthetic viable interactions, wherein an essential gene is rendered non-essential in a gene deletion background^10^, should result in a lack of growth inhibition at all concentrations of inducer tested and should be discernible at high concentrations of inducer – 500 ng/ml aTc. In the case of Lpp and lipoprotein synthesis/transport, genes involved in synthesis/transport remain essential in an Lpp-deficient strain^55^. However, Δ*lpp* bacteria are more tolerant to perturbations in these genes, presumably due to the decreased rate of lipoprotein mis-localization, and accordingly are more resistant to chemical inhibitors of lipoprotein transport and biogenesis^42^. Although screening at the highest inducer concentration alone would have identified the interaction between Lpp and lipoprotein transport, screening at a range of inducer concentrations informed on the global effects of an Δ*lpp* background on the relative essentiality of different processes.

To investigate the inverse genetic interaction screen, where a single CRISPRi construct is conjugated into an existing genomic collection, we chose to conjugate the strongest suppressor from the Δ*lpp*-CRISPRi conjugation screen, the *lolA* CRISPRi knockdown, into all ∼4,000 strains of the Keio collection. High-throughput conjugation of a CRISPRi plasmid into the Keio collection was performed as before, with some minor modifications to the methodology (**Figure S7**). Here, we were able to investigate the effects of disrupting an essential process by CRISPRi in 4,000 non-essential deletion strains simultaneously, wherein 384 distinct deletion strains are found on each screening plate. As each deletion strain in the Keio could potentially grow faster or slower than its neighbors, it was crucial to include an empty vector control for each deletion mutant. As such, we designed an assay setup that permits screening 3 different CRISPRi knockdowns and an empty vector control in 1,536 colony density arrays, such that each mutant harboring the control plasmid is proximal to the same mutant harboring different CRISPRi plasmids (**Figure S7B**).

In addition to conjugating and screening the *lolA* CRISPRi knockdown with the Keio collection, we also conjugated and screened CRISPRi knockdowns of *pssA* and *mreD*, which code for phosphatidylserine synthase^57^ and an integral inner-membrane component of the elongasome^58^, respectively, as controls to identify genetic interactions specific to *lolA*. This is crucial because we have previously observed a ‘frequent hitter’ phenomenon when introducing a second non-essential deletion to the Keio collection with synthetic genetic arrays^21^. Specifically, mutant strains with deficiencies in conjugation or recombination were frequently found to be enhancers or suppressors of growth. With CRISPRi Keio experiments, ‘frequent hitters’ are likely to be deletion mutants that have deficiencies in conjugation, or impact an aspect of the mechanism by which CRISPRi silences gene expression. This includes deletion mutants with altered permeability or accumulation of the inducer (aTc), or more interestingly, mutants that affect the mechanism of gene silencing by CRISPR. This could provide valuable insight into the underlying biology of CRISPRi mediated gene silencing in bacteria and possible routes for a bacterium to acquire resistance to CRISPRi mediated growth inhibition.

The CRISPRi-Keio screen was conducted at two concentrations of inducer, 50 and 500 ng/ml aTc (**Supplementary Table 4**). We hypothesized that enhancers of growth inhibition by CRISPRi would be easier to identify at lower concentration of inducers with marginal levels of inhibition, whilst suppressors would be apparent at higher concentrations of inducer and more severe levels of inhibition. Indeed, with increasing concentrations of inducer we observed that mutants harboring the empty vectors were unperturbed whereas mutants harboring essential gene knockdowns exhibited a dose dependent growth defect across the screening plate (**Figure 2D, Figure S8**). To normalize the growth of each colony, we employed a previously established method that accounts for both the positional effects of a colony and plate-to-plate variability observed in high-density screening approaches^59^ (**Figure S9**), described in detail within the methods section. After normalization, growth of each deletion mutant with a knockdown plasmid was then compared to the growth of that respective deletion mutant harboring the empty vector.

Using a cut-off of 3-standard deviations from the mean of the dataset, we identified 68 deletion mutants that suppressed growth inhibition from *lolA* knockdown at 500 ng/ml aTc. As expected, we found that the Δ*lpp* mutant was one of the strongest suppressors **(Figure 2E)**. To a lesser degree, we also identified Δ*fis* as a suppressor of growth inhibition by *lolA* knockdown. Fis deficient *E. coli* has been shown to be resistant to growth defects associated with an accumulation of a mutant Lpp which is unable to proceed through the lipoprotein processing and transport^60^, similar to the stress imposed by knocking down *lolA* expression by CRISPRi. Interestingly, we also identified gene deletions that suppressed growth inhibition to a similar degree as Δ*lpp:* Δ*uup*, Δ*cheY* and Δ*holC*. CheY has been shown to act as a flagellar response regulator^61^; while both Uup and HolC have been implicated in aiding stalled replication complexes by associating with the Holliday junctions of stalled replication forks^62^ or by resolving toxic conflicts between the replication and transcription complexes^63^, respectively. Interestingly, of the three mutants, Δ*uup* also suppressed growth inhibition by *mreD* knockdown, and Δ*holC* was likewise a suppressor of growth inhibition by *pssA* knockdown (**Figure S10B**). Neither of these mutants, however, were suppressors across all three Keio crosses. Further investigation is required to test whether these deletion strains are promiscuous suppressors of CRISPRi. In contrast to the many suppressors identified, we found just nine deletion strains showing enhanced growth inhibition from *lolA* knockdown at 50 ng/ml aTc. Of these nine enhancers, two deletion backgrounds showed increased growth inhibition for all three CRISPRi constructs screened, namely Δ*galU* and Δ*yiiS*. It is likely that both deletion strains enhance growth inhibition by potentiating an innate aspect of the CRISPRi system used in our study. A defect in *galU* has been associated with disrupted cell envelope biosynthesis including that of lipopolysaccharide^64,65^, whilst *E. coli* deficient in the σ^E^ regulated *yiiS* has been found to have defects in envelope biogenesis^66,67^. As such, it is likely these mutants have enhanced growth inhibition by CRISPRi due to an increased permeability or accumulation of the inducer compound, aTc.

Herein, we present a new tool and methodology to study genetic interactions of the *E. coli* essential genome that is compatible with existing genomic collections. Notably, the approach is translatable to other bacterial pathogens with establish ordered genomic collections. pFD152 and sister plasmids described by Depardieu and Bikard^40^ are built on a broad host-range plasmid compatible with *Salmonella* Typhimurium, *Acinetobacter baumannii, Klebsiella pneumoniae* and *Staphylococcus aureus* amongst other bacteria. Analogous collections of CRISPRi plasmids could be constructed on this backbone and used as tools to investigate genetic interactions in these other important human pathogens. The *E. coli* CRISPRi collection described here, can be used to investigate the dispensability of essential processes in different genetic and environmental contexts and to enhance our understanding of essential gene biology, as well as the downstream effects of perturbing different biological processes. This information could then be used to design unconventional screening approaches to identify chemical inhibitors of previously untargeted processes, or identify growth situations where problematic pathogens cannot survive.

## Acknowledgments

We would like to thank Drs. Shawn French and Maya Farha for helpful comments and discussion. This research was supported by a Discovery Grant for the Natural Sciences and Engineering Research Council of Canada (RGPIN-2019-07090) and by infrastructure funding from Canada Foundation for Innovation and the Ontario Research Fund (ORF-RE09-047). E.D.B was supported by a tier I Canada research chair award, K.R. was supported by an Ontario graduate scholarship, and M.M.T. was supported by Canada Graduate Scholarship.

## Author Contributions

Conceptualization, K.R. and E.D.B.; Methodology, K.R. and E.D.B.; Investigation, K.R., M.M.T., S.J.M., and D.H.; Formal Analysis, K.R.; Writing – Original Draft, K.R, and E.D.B.; Writing – Review & Editing, M.M.T., and E.D.B.; Funding Acquisition, E.D.B; Resources, E.D.B; Supervision, E.D.B.

## Methods

### Growth conditions, plasmids, and strains

Bacteria were routinely cultured in LB medium at 37°C (10 g/L NaCl, 10 g/L tryptone, 5 g/L yeast extract), supplemented with antibiotics (kanamycin, 50 μg/mL and spectinomycin, 150 μg/mL) or 10 mM Diaminopimelic acid (Sigma Aldrich, cat# D1377) when required. For experiments on solid medium, media was prepared with 15 g/L agar. MOPS minimal medium (Teknova) was prepared according to the manufacturer’s instructions: components were filter sterilized after preparing liquid growth medium or added to sterile water and agar (1.5% wt/vol) for solid growth medium. Anhydrotetracycline (aTc) was added to solid agar media to induce CRISPRi when necessary.

sgRNAs targeting different essential genes in *E. coli* were cloned into the conjugative CRISPRi plasmid pFD152 using a previously described single step golden gate assembly protocol^40^. pFD152 was a gift from David Bikard (Addgene plasmid # 125546). sgRNAs used in this study were designed using the publicly available CRISPRbact tool (https://gitlab.pasteur.fr/dbikard/crisprbact), wherein, an sgRNA with the highest predicted on-target activity score and the least off-target homology was selected for each gene.

Oligonucleotides for cloning sgRNAs were ordered from Sigma Aldrich, and specific sequences of oligonucleotides and sequences of sgRNAs can be found in **Supplemental Table 1**. All plasmids constructed were verified by Sanger sequencing before being used for downstream experiments.

The mobile arrayed CRISPRi collection was constructed by transforming sequence verified plasmids into *E. coli* MFD*pir*^43^, the conjugative donor strain for high-throughput conjugation experiments which is auxotrophic for diaminopimelic acid (DAP). All fitness screens with the CRISPRi collection were performed in *E. coli* 25113 [F-Δ*(araD-araB*)567 *lacZ*4787Δ*::rrnB*-3 LAM-rph-1 Δ*(rhaD-rhaB*)568 hsdR514], and, all genes deletions used in the study were obtained from the Keio collection^4^ (kanamycin resistant single-gene deletions in *E. coli* BW25113).

### High-Throughput Conjugation and CRISPRi Array Screening

High-throughput conjugation of the mobile arrayed CRISPRi collection was conducted as previously described with some modifications^20–22^. For high-throughput conjugation and screening of the CRISPRi collection into a genetic background of interest such as *E. coli* Δ*lpp*, the query strain (kanamycin resistant *E. coli* Δ*lpp*) was arrayed on LB agar with kanamycin at 1,536-colony density using the Singer Rotor HDA. The mobile arrayed CRISPRi collection was likewise pinned at 1,536 colony density, with each strain of the collection in technical quadruplicate, on LB with spectinomycin and DAP. The query strain and CRISPRi collection were then co-pinned onto LB agar with DAP but without antibiotic selection, and incubated at 37°C for 2 h for allow for conjugation to occur. After conjugation, colonies were transferred to LB agar with kanamycin and spectinomycin to select for the exconjugants which have both the gene deletion of interest and the CRISPRi plasmid, then were incubated overnight at 37°C. For screening the CRISPRi collection, the exconjugants were used to inoculate LB media with kanamycin and spectinomycin in a 384-microwell plate (Corning) and grown overnight, then cultures were diluted either 100-fold of 10-fold into sterile PBS for assays in rich or minimal media assays, respectively. Cells were then pinned onto solid media containing six concentrations of aTc (0, 5, 10, 50, 500 ng/ml) using the Singer Rotor HDA, then were grown for 16 hours at 37°C and visualized using transmissive scanners.

For conjugation of a specific CRISPRi plasmid into a genomic collection, the workflow is followed as above with minor modifications. *E. coli* MFD*pir* harboring a CRISPRi plasmid of interest was arrayed at 1,536 density on LB with spectinomycin and DAP, whilst the Keio collection was arrayed at 1,536 density on LB with kanamycin. As before, the donor strain harboring the CRISPRi plasmid and the Keio collection were co-pinned onto LB agar with DAP but without antibiotic selection, and incubated at 37°C for 2 h for allow for conjugation to occur. Then, colonies were transferred to LB agar with kanamycin and spectinomycin to select for the exconjugants which have both the gene deletion of interest and the CRISPRi plasmid, then were incubated overnight at 37°C. For screening the CRISPRi-Keio collections, the exconjugants containing the empty vector and CRISPRi plasmids were first used to inoculate LB with spectinomycin and kanamycin in 384-microwell plates and grown overnight–the Keio collection is 12 individual 384 well plates in its entirety. Then, cultures are spotted onto a 1536-colony density solid agar plates containing different concentrations of aTc, such that each Keio mutant is found in four proximal colonies– each with a different CRISPRi plasmid or vector control. CRISPRi Keio screens were conducted in technical duplicate.

### Plate imaging quantification and analysis

After being grown to endpoint, solid media screening plates were imaged using an Epson Perfection V750 to generate a high-resolution image, from which the integrated density of each colony was determined using previously published ImageJ pipelines^12^. Analysis and data normalization was performed differently based on screen being performed. For analysis of CRISPRi collection growth in wildtype (*lpp*+) and Δ*lpp* backgrounds, the average raw colony density of the 17 strains harboring the empty vector was calculated, then, this value was used to normalize the growth of strains harboring different CRISPRi plasmids. The average of the two technical replicates was then calculated and is reported in Supplemental Table 2. As the whole collection was screened on a single solid agar plate, plate to plate variability was not a concern, and spatial growth effects likewise did not appear to impact growth meaningfully.

For analysis of the CRISPRi Keio assay, a two-pass normalization method was used to account for both spatial and plate to plate variation observed. This normalization is similar to a previously published approach to normalize high-throughput screening datasets^59^. In summary, the entire Keio screen used 24 assay plates at every aTc concentration – the 12 Keio assay plates screened in biological duplicate – wherein mutants with the same CRISPRi plasmid occupy the same position in each plate. First, spatial effects across the 24 assay plates were normalized by dividing each colony with the inter-quartile mean (IQM) of every colony occupying the same position on other plates (with the same CRISPRi plasmid). Then, plate to plate variability was normalized by dividing the growth of each colony, by the IQM of the growth of every other colony harboring the same CRISPRi plasmid within each screening plate. Following spatial and intra-plate normalization, the growth of each deletion mutant with a gene knockdown plasmid was then normalized to the growth that respective deletion mutant with the empty vector.

### MIC determination and dilution plating

For MIC assays, overnight cultures of bacteria were grown in LB broth supplemented with appropriate antibiotic selection. Cultures were diluted (1:100) into fresh LB media and grown to mid-log phase of growth (OD ∼0.5). Subcultures were diluted to an OD of 0.1, then diluted (1:5000) into assay media, MOPS minimal media or LB. MIC assay plates were incubated at 37°C for 18 hours, then OD_600_ was measured using a Tecan Infinite series micro-plate reader. For dilution plating assays, overnight cultures of bacteria were grown in LB broth supplemented with appropriate antibiotic selection. Following growth, each culture was pelleted and resuspended in sterile PBS to a final OD of 1.0. Then 10-fold serial dilutions of the culture were prepared in sterile PBS, and a 10 μl spot of dispensed onto solid growth plates with and without aTc, and grown for 18 hours at 37°C. Images of dilution plating assay plates were obtained using an Epson Perfection V750 high-resolution scanner.

## References

1. Hogan, A.M., and Cardona, S.T. (2022). Gradients in gene essentiality reshape antibacterial research. FEMS Microbiol Rev 46.p 10.1093/FEMSRE/FUAC005.

2. D’Elia, M.A., Pereira, M.P., and Brown, E.D. (2009). Are essential genes really essential? Trends Microbiol 17, 433–438. 10.1016/j.tim.2009.08.005.

3. Goodall, E.C.A., Robinson, A., Johnston, I.G., Jabbari, S., Turner, K.A., Cunningham, A.F., Lund, P.A., Cole, J.A., and Henderson, I.R. (2018). The essential genome of Escherichia coli K-12. mBio 9, e02096–17. 10.1128/mBio.02096-17.

4. Baba, T., Ara, T., Hasegawa, M., Takai, Y., Okumura, Y., Baba, M., Datsenko, K.A., Tomita, M., Wanner, B.L., and Mori, H. (2006). Construction of Escherichia coli K-12 in-frame, single-gene knockout mutants: the Keio collection. Mol Syst Biol 2, 2006–0008. 10.1038/msb4100050.

5. Rousset, F., Cui, L., Siouve, E., Becavin, C., Depardieu, F., and Bikard, D. (2018). Genomewide CRISPR-dCas9 screens in E. coli identify essential genes and phage host factors. PLoS Genet 14, e1007749. 10.1371/JOURNAL.PGEN.1007749.

6. Gerdes, S.Y., Scholle, M.D., Campbell, J.W., Balázsi, G., Ravasz, E., Daugherty, M.D., Somera, A.L., Kyrpides, N.C., Anderson, I., Gelfand, M.S., et al. (2003). Experimental determination and system level analysis of essential genes in Escherichia coli MG1655. J Bacteriol 185, 5673–5684. 10.1128/JB.185.19.5673-5684.2003.

7. Juhas, M., Eberl, L., and Church, G.M. (2012). Essential genes as antimicrobial targets and cornerstones of synthetic biology. Trends Biotechnol 30, 601–607. 10.1016/J.TIBTECH.2012.08.002.

8. Farha, M.A., Leung, A., Sewell, E.W., D’Elia, M.A., Allison, S.E., Ejim, L., Pereira, P.M., Pinho, M.G., Wright, G.D., and Brown, E.D. (2013). Inhibition of WTA synthesis blocks the cooperative action of pbps and sensitizes MRSA to β-lactams. ACS Chem Biol 8, 226–233. 10.1021/cb300413m.

9. Gehrke, S.S., Kumar, G., Yokubynas, N.A., Côté, J.-P., Wang, W., French, S., MacNair, C.R., Wright, G.D., and Brown, E.D. (2017). Exploiting the Sensitivity of Nutrient Transporter Deletion Strains in Discovery of Natural Product Antimetabolites. ACS Infect Dis 3, 955– 965. 10.1021/ACSINFECDIS.7B00149.

10. Klobucar, K., and Brown, E.D. (2018). Use of genetic and chemical synthetic lethality as probes of complexity in bacterial cell systems. FEMS Microbiol Rev 42, fux054. 10.1093/femsre/fux054.

11. Nichols, R.J., Sen, S., Choo, Y.J., Beltrao, P., Zietek, M., Chaba, R., Lee, S., Kazmierczak, K.M., Lee, K.J., Wong, A., et al. (2011). Phenotypic landscape of a bacterial cell. Cell 144, 143–156. 10.1016/j.cell.2010.11.052.

12. French, S., Mangat, C., Bharat, A., Cté, J.P., Mori, H., and Brown, E.D. (2016). A robust platform for chemical genomics in bacterial systems. Mol Biol Cell 27, 1015–1025. 10.1091/mbc.E15-08-0573.

13. Tong, M., French, S., El Zahed, S.S., Ong, W. kit Karp, P.D., and Brown, E.D. (2020). Gene Dispensability in Escherichia coli Grown in Thirty Different Carbon Environments. mBio 11, e02259–20. 10.1128/mBio.02259-20.

14. Typas, A., Nichols, R.J., Siegele, D.A., Shales, M., Collins, S.R., Lim, B., Braberg, H., Yamamoto, N., Takeuchi, R., Wanner, B.L., et al. (2008). High-throughput, quantitative analyses of genetic interactions in E. coli. Nat Methods 5, 781–787. 10.1038/nmeth.1240.

15. Babu, M., Díaz-Mejía, J.J., Vlasblom, J., Gagarinova, A., Phanse, S., Graham, C., Yousif, F., Ding, H., Xiong, X., Nazarians-Armavil, A., et al. (2011). Genetic interaction maps in Escherichia coli reveal functional crosstalk among cell envelope biogenesis pathways. PLoS Genet 7, e1002377. 10.1371/journal.pgen.1002377.

16. Babu, M., Arnold, R., Bundalovic-Torma, C., Gagarinova, A., Wong, K.S., Kumar, A., Stewart, G., Samanfar, B., Aoki, H., Wagih, O., et al. (2014). Quantitative Genome-Wide Genetic Interaction Screens Reveal Global Epistatic Relationships of Protein Complexes in Escherichia coli. PLoS Genet 10, e1004120. 10.1371/journal.pgen.1004120.

17. Butland, G., Babu, M., Díaz-Mejía, J.J., Bohdana, F., Phanse, S., Gold, B., Yang, W., Li, J., Gagarinova, A.G., Pogoutse, O., et al. (2008). eSGA: E. coli synthetic genetic array analysis. Nat Methods 5, 789–795. 10.1038/nmeth.1239.

18. Babu, M., Bundalovic-Torma, C., Calmettes, C., Phanse, S., Zhang, Q., Jiang, Y., Minic, Z., Kim, S., Mehla, J., Gagarinova, A., et al. (2018). Global landscape of cell envelope protein complexes in Escherichia coli. Nat Biotechnol 13, 103–112. 10.1038/nbt.4024.

19. Kumar, A., Beloglazova, N., Bundalovic-Torma, C., Phanse, S., Deineko, V., Gagarinova, A., Musso, G., Vlasblom, J., Lemak, S., Hooshyar, M., et al. (2016). Conditional Epistatic Interaction Maps Reveal Global Functional Rewiring of Genome Integrity Pathways in Escherichia coli. Cell Rep 14, 648–661. 10.1016/j.celrep.2015.12.060.

20. Rachwalski, K., Ellis, M.J., Tong, M., and Brown, E.D. (2022). Synthetic Genetic Interactions Reveal a Dense and Cryptic Regulatory Network of Small Noncoding RNAs in Escherichia coli. mBio 13, e0122522. 10.1128/MBIO.01225-22.

21. Klobucar, K., French, S., Côté, J.P., Howes, J.R., and Brown, E.D. (2020). Genetic and chemical-genetic interactions map biogenesis and permeability determinants of the outer membrane of Escherichia coli. mBio 11, 1–17. 10.1128/mBio.00161-20.

22. Côté, J.P., French, S., Gehrke, S.S., MacNair, C.R., Mangat, C.S., Bharat, A., and Brown, E.D. (2016). The genome-wide interaction network of nutrient stress genes in Escherichia coli. mBio 7, 1–12. 10.1128/mBio.01714-16.

23. Price, M.N., Wetmore, K.M., Waters, R.J., Callaghan, M., Ray, J., Liu, H., Kuehl, J. V., Melnyk, R.A., Lamson, J.S., Suh, Y., et al. (2018). Mutant phenotypes for thousands of bacterial genes of unknown function. Nature 557, 503–509. 10.1038/s41586-018-0124-0.

24. French, S., Ellis, M.J., Coutts, B.E., and Brown, E.D. (2017). Chemical genomics reveals mechanistic hypotheses for uncharacterized bioactive molecules in bacteria. Curr Opin Microbiol 39, 42–47. 10.1016/J.MIB.2017.09.005.

25. Larson, M.H., Gilbert, L.A., Wang, X., Lim, W.A., Weissman, J.S., and Qi, L.S. (2013). CRISPR interference (CRISPRi) for sequence-specific control of gene expression. Nat Protoc 8, 2180–2196. 10.1038/nprot.2013.132.

26. Todor, H., Silvis, M.R., Osadnik, H., and Gross, C.A. (2021). Bacterial CRISPR screens for gene function. Curr Opin Microbiol 59, 102–109. 10.1016/J.MIB.2020.11.005.

27. Peters, J.M., Koo, B.-M., Patino, R., Heussler, G.E., Hearne, C.C., Qu, J., Inclan, Y.F., Hawkins, J.S., Lu, C.H.S., Silvis, M.R., et al. (2019). Enabling genetic analysis of diverse bacteria with Mobile-CRISPRi. Nature Microbiology 2019 4:2 4, 244–250. 10.1038/s41564-018-0327-z.

28. Keseler, I.M., Gama-Castro, S., Mackie, A., Billington, R., Bonavides-Martínez, C., Caspi, R., Kothari, A., Krummenacker, M., Midford, P.E., Muñiz-Rascado, L., et al. (2021). The EcoCyc Database in 2021. Front Microbiol 12, 2098. 10.3389/FMICB.2021.711077.

29. Li, S., Jendresen, C.B., Landberg, J., Pedersen, L.E., Sonnenschein, N., Jensen, S.I., and Nielsen, A.T. (2020). Genome-Wide CRISPRi-Based Identification of Targets for Decoupling Growth from Production. ACS Synth Biol 9, 1030–1040. 10.1021/ACSSYNBIO.9B00143.

30. Cui, L., Vigouroux, A., Rousset, F., Varet, H., Khanna, V., and Bikard, D. (2018). A CRISPRi screen in E. coli reveals sequence-specific toxicity of dCas9. Nat Commun 9, 1912. 10.1038/s41467-018-04209-5.

31. Calvo-Villamã, A., Wong Ng, J., Planel, R., Ménager, H., Chen, A., Cui, L., and Bikard, D. (2020). On-target activity predictions enable improved CRISPR-dCas9 screens in bacteria. Nucleic Acids Res 48, 64. 10.1093/nar/gkaa294.

32. de Bakker, V., Liu, X., Bravo, A.M., and Veening, J.W. (2022). CRISPRi-seq for genomewide fitness quantification in bacteria. Nat Protoc 17, 252–281. 10.1038/s41596-021-00639-6.

33. Ranava, D., Yang, Y., Orenday-Tapia, L., Rousset, F., Turlan, C., Morales, V., Cui, L., Moulin, C., Froment, C., Munoz, G., et al. (2021). Lipoprotein dolP supports proper folding of bamA in the bacterial outer membrane promoting fitness upon envelope stress. Elife 10, e67817. 10.7554/ELIFE.67817.

34. Barrick, J.E., and Lenski, R.E. (2009). Genome-wide Mutational Diversity in an Evolving Population of Escherichia coli. Cold Spring Harb Symp Quant Biol 74, sqb.2009.74.018. 10.1101/SQB.2009.74.018.

35. Silvis, M.R., Rajendram, M., Shi, H., Osadnik, H., Gray, A.N., Cesar, S., Peters, J.M., Hearne, C.C., Kumar, P., Todor, H., et al. (2021). Morphological and transcriptional responses to CRISPRi knockdown of essential genes in Escherichia coli. mBio 12, e0256121. 10.1128/MBIO.02561-21.

36. Peters, J.M., Colavin, A., Shi, H., Czarny, T.L., Larson, M.H., Wong, S., Hawkins, J.S., Lu, C.H.S., Koo, B.M., Marta, E., et al. (2016). A comprehensive, CRISPR-based functional analysis of essential genes in bacteria. Cell 165, 1493–1506. 10.1016/j.cell.2016.05.003.

37. Anglada-Girotto, M., Handschin, G., Ortmayr, K., Campos, A.I., Gillet, L., Manfredi, P., Mulholland, C. V., Berney, M., Jenal, U., Picotti, P., et al. (2022). Combining CRISPRi and metabolomics for functional annotation of compound libraries. Nature Chemical Biology 2022 18:5 18, 482–491. 10.1038/s41589-022-00970-3.

38. Donati, S., Kuntz, M., Pahl, V., Farke, N., Beuter, D., Glatter, T., Gomes-Filho, J.V., Randau, L., Wang, C.Y., and Link, H. (2021). Multi-omics Analysis of CRISPRi-Knockdowns Identifies Mechanisms that Buffer Decreases of Enzymes in E. coli Metabolism. Cell Syst 12, 56–67.e6. 10.1016/J.CELS.2020.10.011.

39. Mateus, A., Hevler, J., Bobonis, J., Kurzawa, N., Shah, M., Mitosch, K., Goemans, C. V., Helm, D., Stein, F., Typas, A., et al. (2020). The functional proteome landscape of Escherichia coli. Nature 588, 473–478. 10.1038/s41586-020-3002-5.

40. Depardieu, F., and Bikard, D. (2019). Gene silencing with CRISPRi in bacteria and optimization of dCas9 expression levels. Methods, 1–15. 10.1016/j.ymeth.2019.07.024.

41. Zwiebel, L.J., Inukai, M., Nakamura, K., and Inouye, M. (1981). Preferential selection of deletion mutations of the outer membrane lipoprotein gene of Escherichia coli by globomycin. J Bacteriol 145, 654–656. 10.1128/JB.145.1.654-656.1981.

42. Lehman, K.M., Smith, H.C., and Grabowicz, M. (2022). A Biological Signature for the Inhibition of Outer Membrane Lipoprotein Biogenesis. mBio 13, e0075722. 10.1128/MBIO.00757-22.

43. Ferrières, L., Hémery, G., Nham, T., Guérout, A.M., Mazel, D., Beloin, C., and Ghigo, J.M. (2010). Silent mischief: Bacteriophage Mu insertions contaminate products of Escherichia coli random mutagenesis performed using suicidal transposon delivery plasmids mobilized by broad-host-range RP4 conjugative machinery. J Bacteriol 192, 6418–6427. 10.1128/JB.00621-10.

44. Zaslaver, A., Bren, A., Ronen, M., Itzkovitz, S., Kikoin, I., Shavit, S., Liebermeister, W., Surette, M.G., and Alon, U. (2006). A comprehensive library of fluorescent transcriptional reporters for Escherichia coli. Nat Methods 3, 623–628. 10.1038/nmeth895.

45. Kitagawa, M., Ara, T., Arifuzzaman, M., Ioka-Nakamichi, T., Inamoto, E., Toyonaga, H., and Mori, H. (2005). Complete set of ORF clones of Escherichia coli ASKA library (A complete set of E. coli K-12 ORF archive): unique resources for biological research. DNA Research 12, 291–299. 10.1093/dnares/dsi012.

46. Braun, V. (1973). Molecular organization of the rigid layer and the cell wall of Escherichia coli. J Infect Dis 128, S9–S16. 10.1093/INFDIS/128.

47. Braun, V., and Rehn, K. (1969). Chemical characterization, spatial distribution and function of a lipoprotein (murein-lipoprotein) of the E. coli cell wall. The specific effect of trypsin on the membrane structure. Eur J Biochem 10, 426–438. 10.1111/J.1432-1033.1969.TB00707.X.

48. Inouye, S., Wang, S., and Sezikawa, J. (1977). Amino acid sequence for the peptide extension on the prolipoprotein of the Escherichia coli outer membrane. Proc Natl Acad Sci U S A 74, 1004–1008. 10.1073/PNAS.74.3.1004.

49. Mathelié-Guinlet, M., Asmar, A.T., Collet, J.F., and Dufrêne, Y.F. (2020). Lipoprotein Lpp regulates the mechanical properties of the E. coli cell envelope. Nature Communications 2020 11:1 11, 1–11. 10.1038/s41467-020-15489-1.

50. Grabowicz, M. (2019). Lipoproteins and Their Trafficking to the Outer Membrane. EcoSal Plus 8. 10.1128/ECOSALPLUS.ESP-0038-2018.

51. Ruiz, N., Gronenberg, L.S., Kahne, D., and Silhavy, T.J. (2008). Identification of two innermembrane proteins required for the transport of lipopolysaccharide to the outer membrane of Escherichia coli. Proc Natl Acad Sci U S A 105, 5537–5542. 10.1073/pnas.2218473120.

52. Grabowicz, M., and Silhavy, T.J. (2017). Redefining the essential trafficking pathway for outer membrane lipoproteins. Proc Natl Acad Sci U S A 114, 4769–4774. 10.1073/PNAS.1702248114.

53. McLeod, S.M., Fleming, P.R., MacCormack, K., McLaughlin, R.E., Whiteaker, J.D., Narita, S. ichiro Mori, M., Tokuda, H., and Miller, A.A. (2015). Small-molecule inhibitors of gram-negative lipoprotein trafficking discovered by phenotypic screening. J Bacteriol 197, 1075–1082. 10.1128/JB.02352-14.

54. Hart, E.M., Mitchell, A.M., Konovalova, A., Grabowicz, M., Sheng, J., Han, X., Rodriguez-Rivera, F.P., Schwaid, A.G., Malinverni, J.C., Balibar, C.J., et al. (2019). A small-molecule inhibitor of BamA impervious to efflux and the outer membrane permeability barrier. PNAS 116, 21748–21757. 10.1073/PNAS.1912345116.

55. Tajima, T., Yokota, N., Matsuyama, S.I., and Tokuda, H. (1998). Genetic analyses of the in vivo function of LolA, a periplasmic chaperone involved in the outer membrane localization of Escherichia coli lipoproteins. FEBS Lett 439, 51–54. 10.1016/S0014-5793(98)01334-9.

56. Diao, J., Komura, R., Sano, T., Pantua, H., Storek, K.M., Inaba, H., Ogawa, H., Noland, C.L., Peng, Y., Gloor, S.L., et al. (2021). Inhibition of Escherichia coli lipoprotein diacylglyceryl transferase is insensitive to resistance caused by deletion of Braun’s lipoprotein. J Bacteriol 203.p 10.1128/JB.00149-21.

57. Raetz, C.R., Carman, G.M., Dowhan, W., Jiang, R.T., Waszkuc, W., Loffredo, W., and Tsa, M.D. (1987). Phospholipids chiral at phosphorus. Steric course of the reactions catalyzed by phosphatidylserine synthase from Escherichia coli and yeast. Biochemistry 26, 4022– 4027. 10.1021/BI00387A042.

58. Liu, X., Biboy, J., Consoli, E., Vollmer, W., and den Blaauwen, T. (2020). MreC and MreD balance the interaction between the elongasome proteins PBP2 and RodA. PLoS Genet 16, e1009276. 10.1371/JOURNAL.PGEN.1009276.

59. Mangat, C.S., Bharat, A., Gehrke, S.S., and Brown, E.D. (2014). Rank Ordering Plate Data Facilitates Data Visualization and Normalization in High-Throughput Screening. J Biomol Screen 19, 1314–1320. 10.1177/1087057114534298.

60. Patil, D., Xun, D., Schueritz, M., Bansal, S., Cheema, A., Crooke, E., and Saxena, R. (2021). Membrane Stress Caused by Unprocessed Outer Membrane Lipoprotein Intermediate Pro-Lpp Affects DnaA and Fis-Dependent Growth. Front Microbiol 12, 1293. 10.3389/FMICB.2021.677812.

61. Antani, J.D., Gupta, R., Lee, A.H., Rhee, K.Y., Manson, M.D., and Lele, P.P. (2021). Mechanosensitive recruitment of stator units promotes binding of the response regulator CheY-P to the flagellar motor. Nat Commun 12, 1–8. 10.1038/s41467-021-25774-2.

62. Romero, Z.J., Armstrong, T.J., Henrikus, S.S., Chen, S.H., Glass, D.J., Ferrazzoli, A.E., Wood, E.A., Chitteni-Pattu, S., Van Oijen, A.M., Lovett, S.T., et al. (2020). Frequent template switching in postreplication gaps: suppression of deleterious consequences by the Escherichia coli Uup and RadD proteins. Nucleic Acids Res 48, 212. 10.1093/NAR/GKZ960.

63. Cooper, D.L., Harada, T., Tamazi, S., Ferrazzoli, A.E., and Lovett, S.T. (2021). The role of replication clamp-loader protein holC of Escherichia coli in overcoming replication/transcription conflicts. mBio 12, 1–15. 10.1128/MBIO.00184-21.

64. Genevaux, P., Bauda, P., DuBow, M.S., and Oudega, B. (1999). Identification of Tn 10 insertions in the rlaG, rfaP, and galU genes involved in lipopolysaccharide core biosynthesis that affect Escherichia coli adhesion. Arch Microbiol 172, 1–8. 10.1007/S002030050732.

65. Sundararajan, T.A., Rapin, A.M.C., and Kalckar, H.M. (1962). Biochemical observations om E. coli mutants defective in uridine diphosphoglucose. PNAS 48, 2187–2193. 10.1073/pnas.48.12.2187.

66. Rezuchova, B., Miticka, H., Homerova, D., Roberts, M., and Kormanec, J. (2003). New members of the Escherichia coli sigmaE regulon identified by a two-plasmid system. FEMS Microbiol Lett 225, 1–7. 10.1016/S0378-1097(03)00480-4.

67. Paradis-Bleau, C., Kritikos, G., Orlova, K., Typas, A., and Bernhardt, T.G. (2014). A Genome-Wide Screen for Bacterial Envelope Biogenesis Mutants Identifies a Novel Factor Involved in Cell Wall Precursor Metabolism. PLoS Genet 10, 1004056. 10.1371/JOURNAL.PGEN.1004056.

