## Supplemental Figures for "A mobile CRISPRi collection enables genetic interaction studies for the essential genes of *Escherichia coli*"

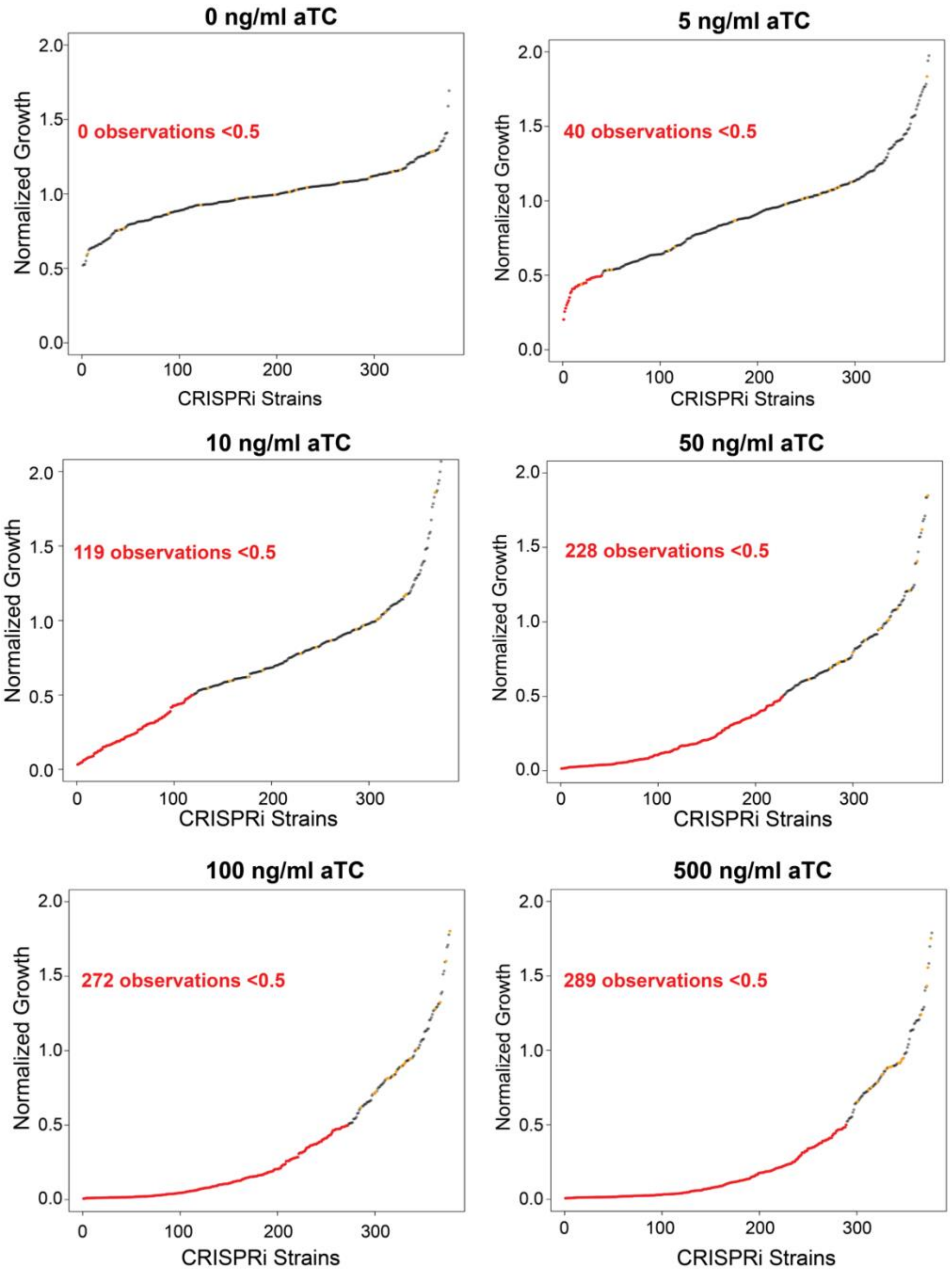


**Figure S1.** Ranked ordered growth of the CRISPRi collection at varying concentrations of aTc on LB media. An overnight culture of the CRISPRi collection was grown in a 384-well plate, then diluted 100-fold into PBS. Bacteria were spotted onto plates containing varying concentrations of aTc using the Singer-ROTOR, and were grown for 16 hours at 37°C. Following incubation, plates were scanned using high-resolution scanners, and integrated colony size was quantified. Strains with growth below 0.5 are shown in red, and the empty vector controls on the plate are shown in orange.

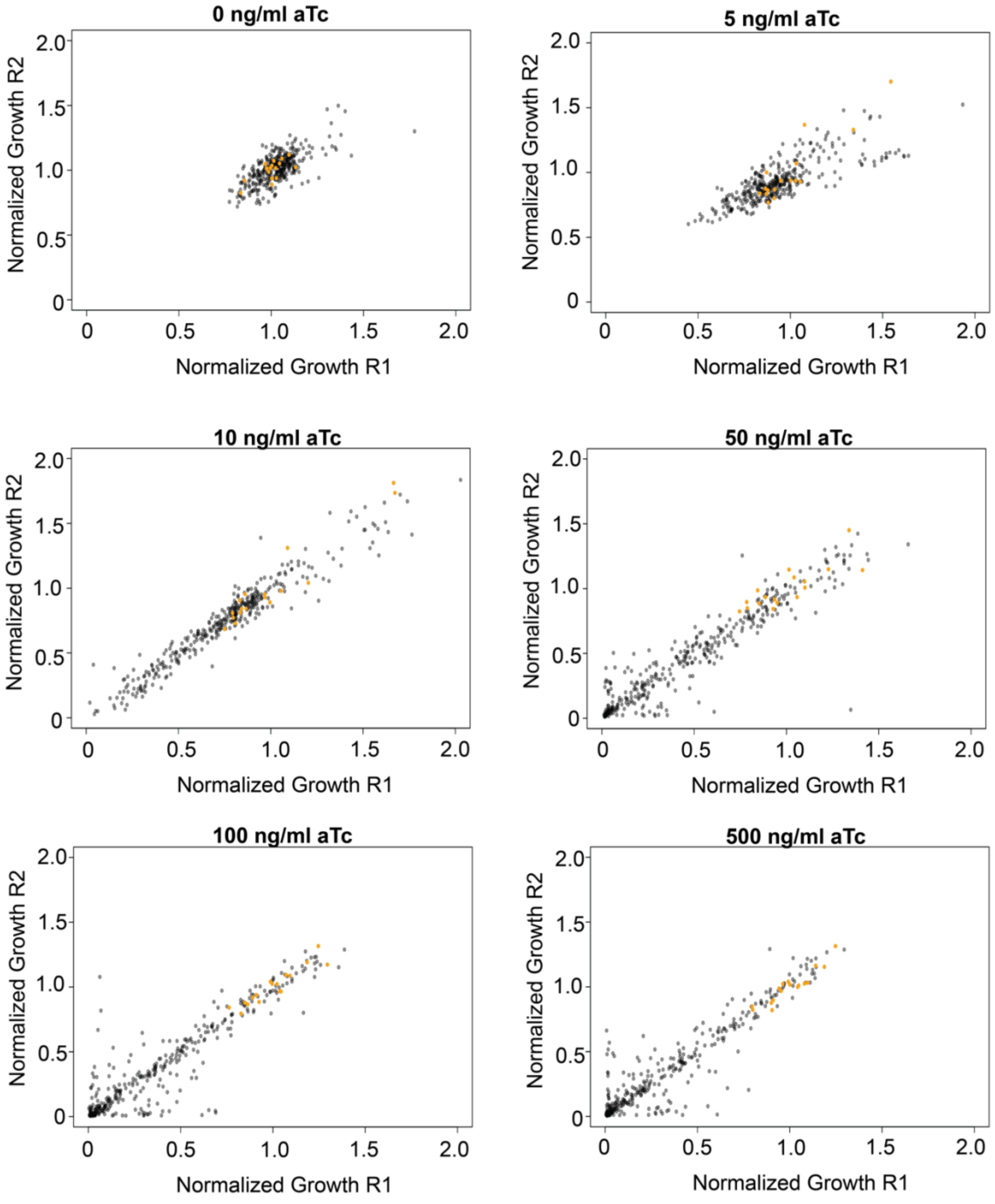


**Figure S2.** Replicate plots of the CRISPRi collection at varying concentrations of aTc on LB media showing good replication across different concentrations of inducer. Empty vector controls on the plate are shown in orange.

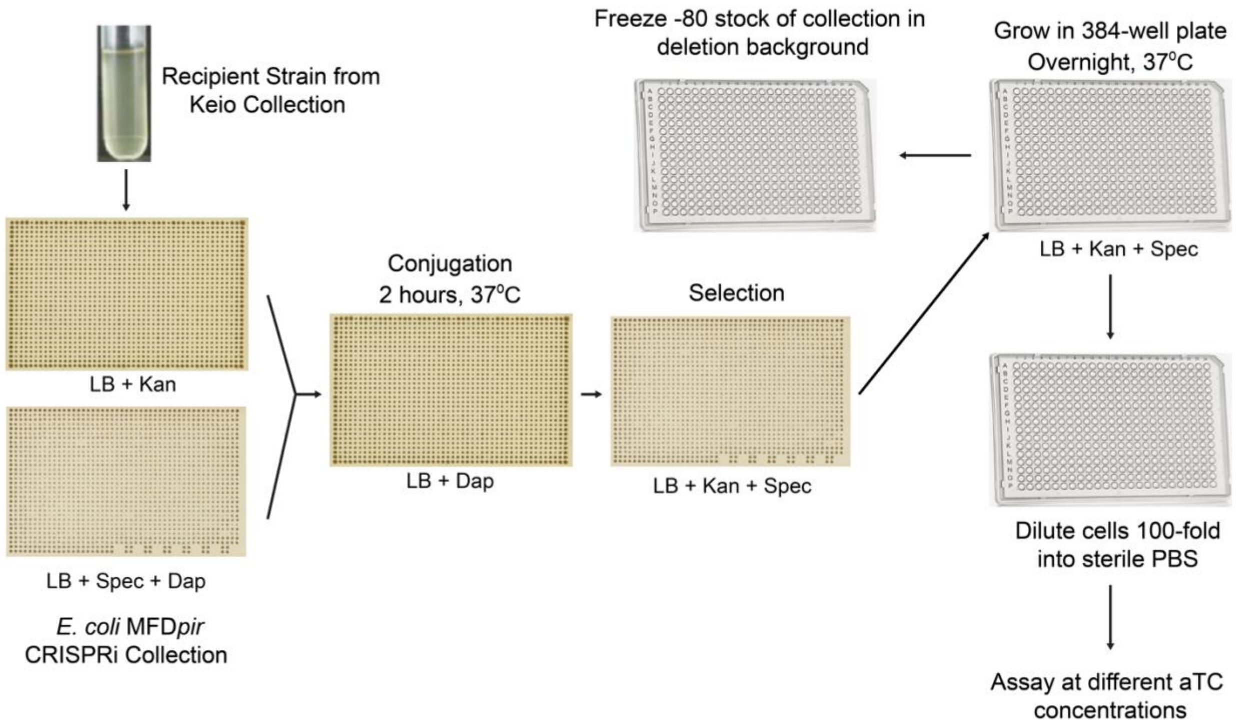


**Figure S3.** Conjugation and assay workflow of the CRISPRi collection into a genetic background of interest. The mobile arrayed CRISPRi collection in *E. coli* MFD*pir* and a recipient strain of interest are co-pinned, and incubated for 2 hours at 37°C, then are pinned onto media containing double selection and grown overnight to select for exconjugants. Exconjugants are then used to inoculate media in a 384-microwell plate and grown overnight. Cells can then be made into a freezer stock for future use, or diluted 100-fold into PBS for assays.

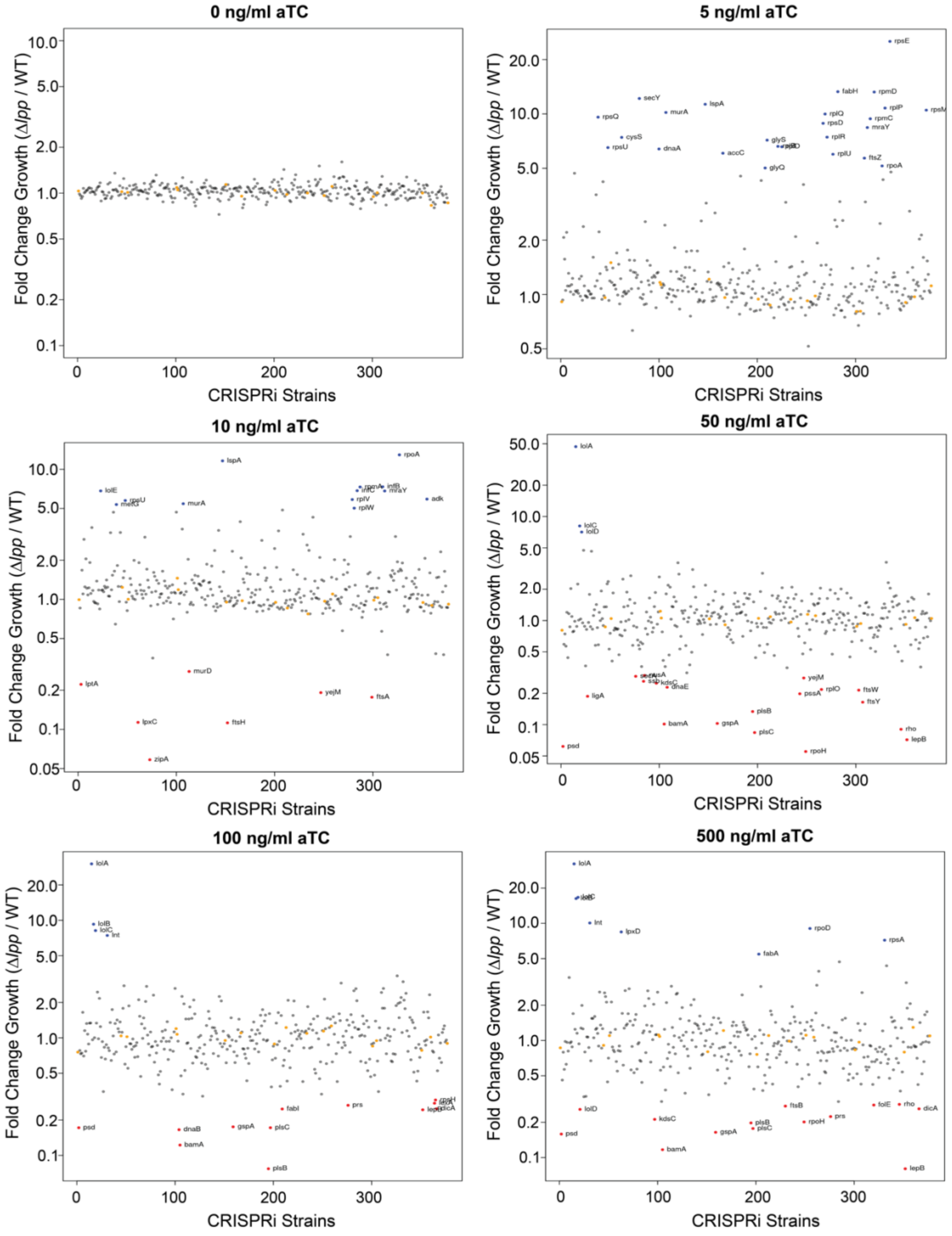


**Figure S4**. Fold-change growth of the Δ*lpp* CRISPRi collection relative to the growth of the WT CRISPRi collection at different concentrations of inducer. Suppressors and enhancers are labelled in blue and red respectively, and were determined as deviating from the mean fold-change of the collection by at least 3-standard deviations. Strains harboring the empty vector are presented in orange.

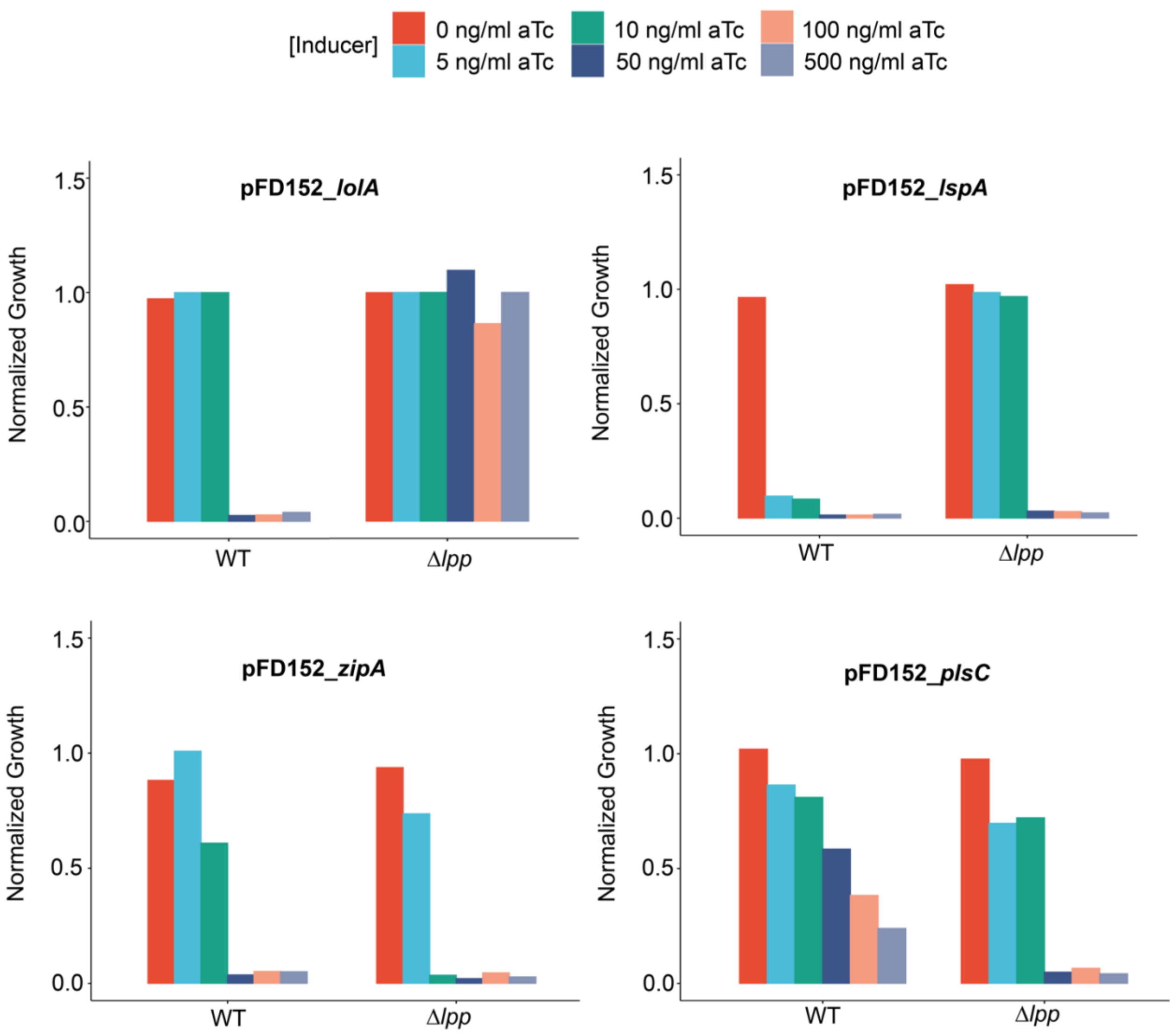


**Figure S5**. Barplots showing the growth of bacteria harboring different CRISPRi plasmids in WT or Δ*lpp* backgrounds at different aTc concentrations.

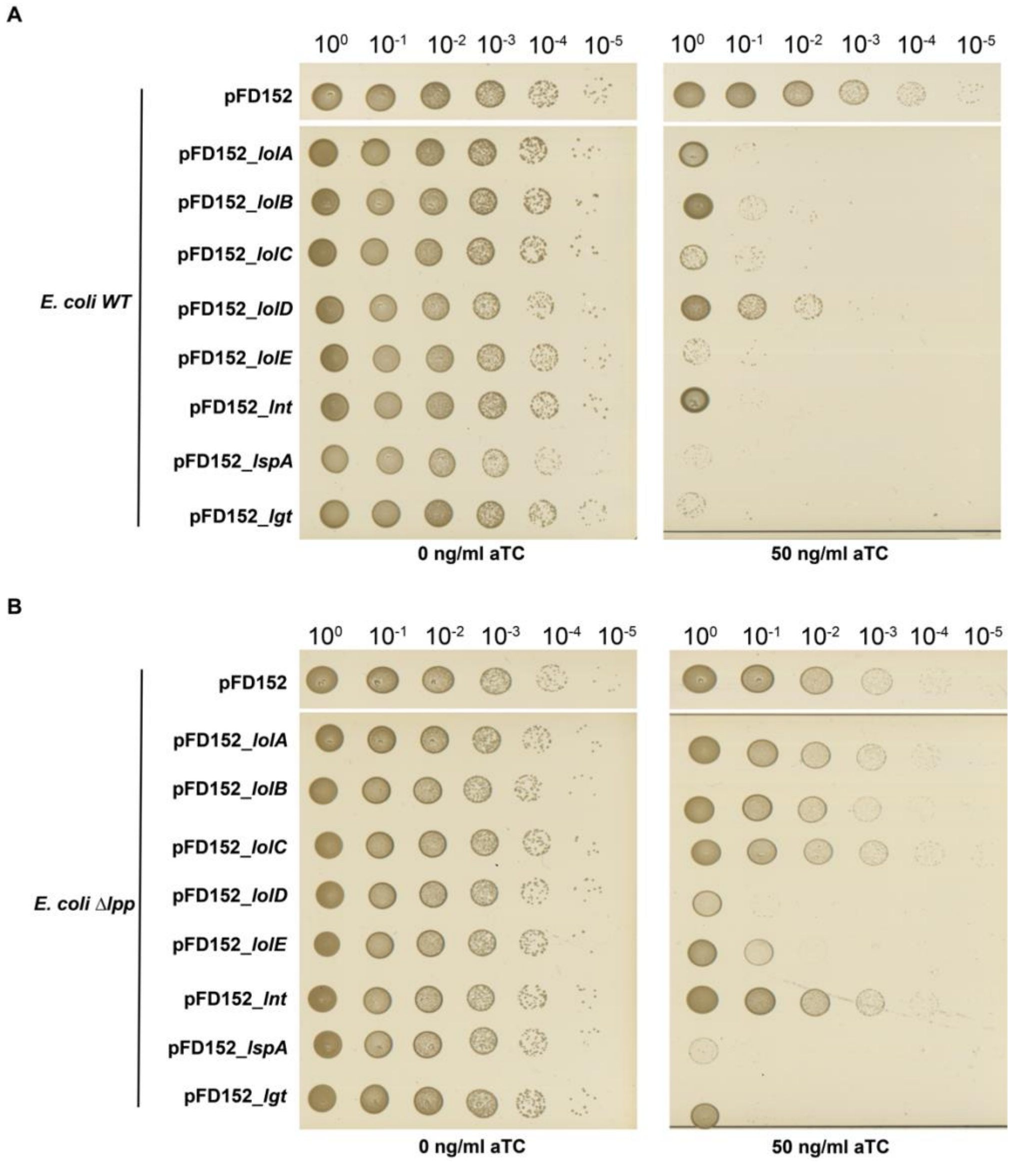


**Figure S6.** Dilution plating *E. coli* containing knockdown plasmids for different genes involved in lipoprotein transport and maturation. 10-fold serial dilutions of overnight cultures from the (A) WT or (B) Δ*lpp* CRISPRi collection were spotted on minimal media containing 0 or 50 ng/ml of aTc to induce CRISPRi. Strong suppression of growth inhibition is seen with knockdown constructs targeting *lolA, lolB, lolC, and lnt*, which were the strongest suppressors of killing in the Δ*lpp* CRISPRi screen.

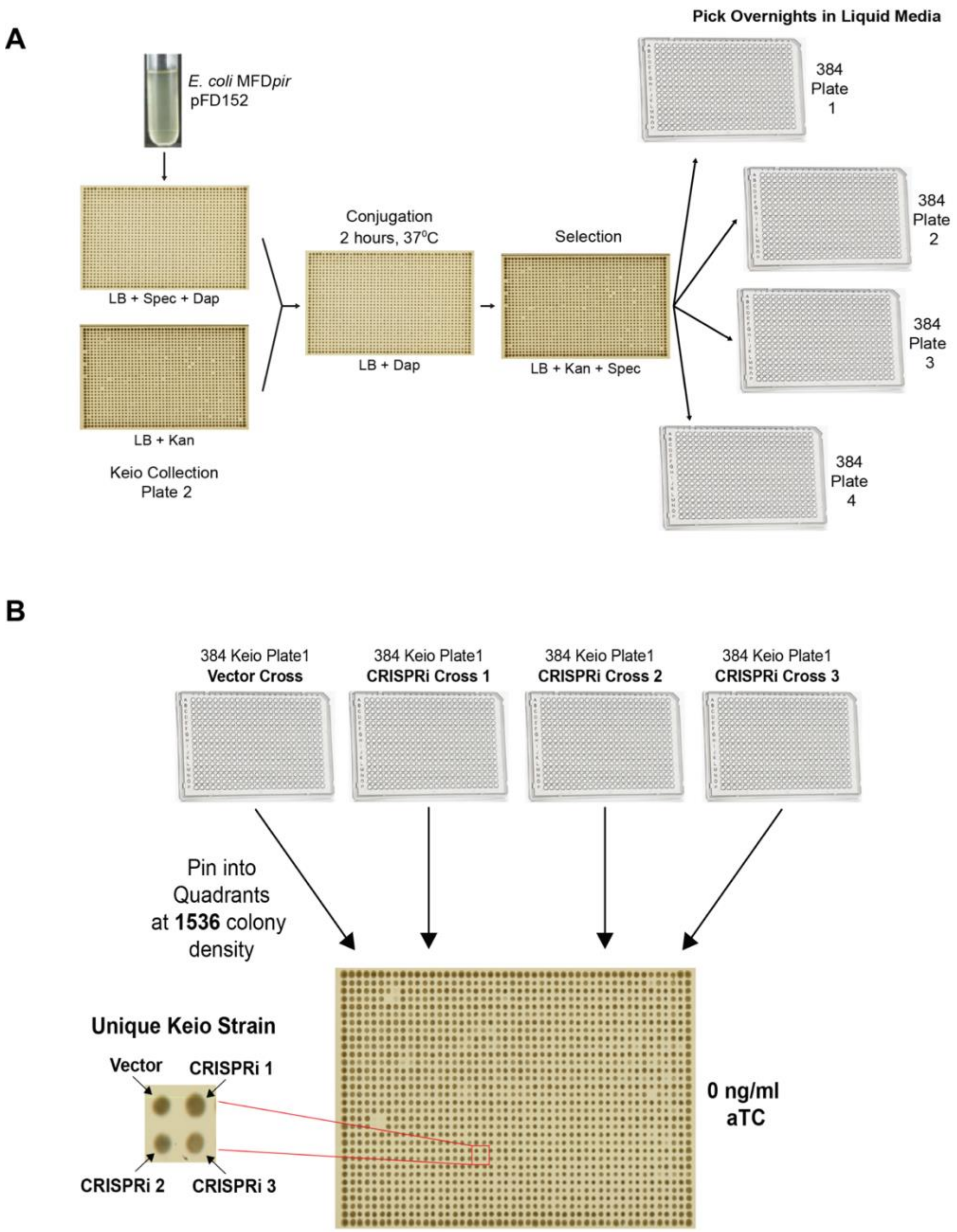


**Figure S7. (A)** Conjugation workflow for introducing CRISPRi constructs into the Keio collection. Keio collection harbouring plasmid was then pinned into media in 384-well plates, and plates grown overnight. **(B)** The Keio collection was screened at 1536 colony density, with each quadrant containing a single deletion strain harbouring 4 different plasmids: an empty vector and 3 different CRISPRi  constructs.

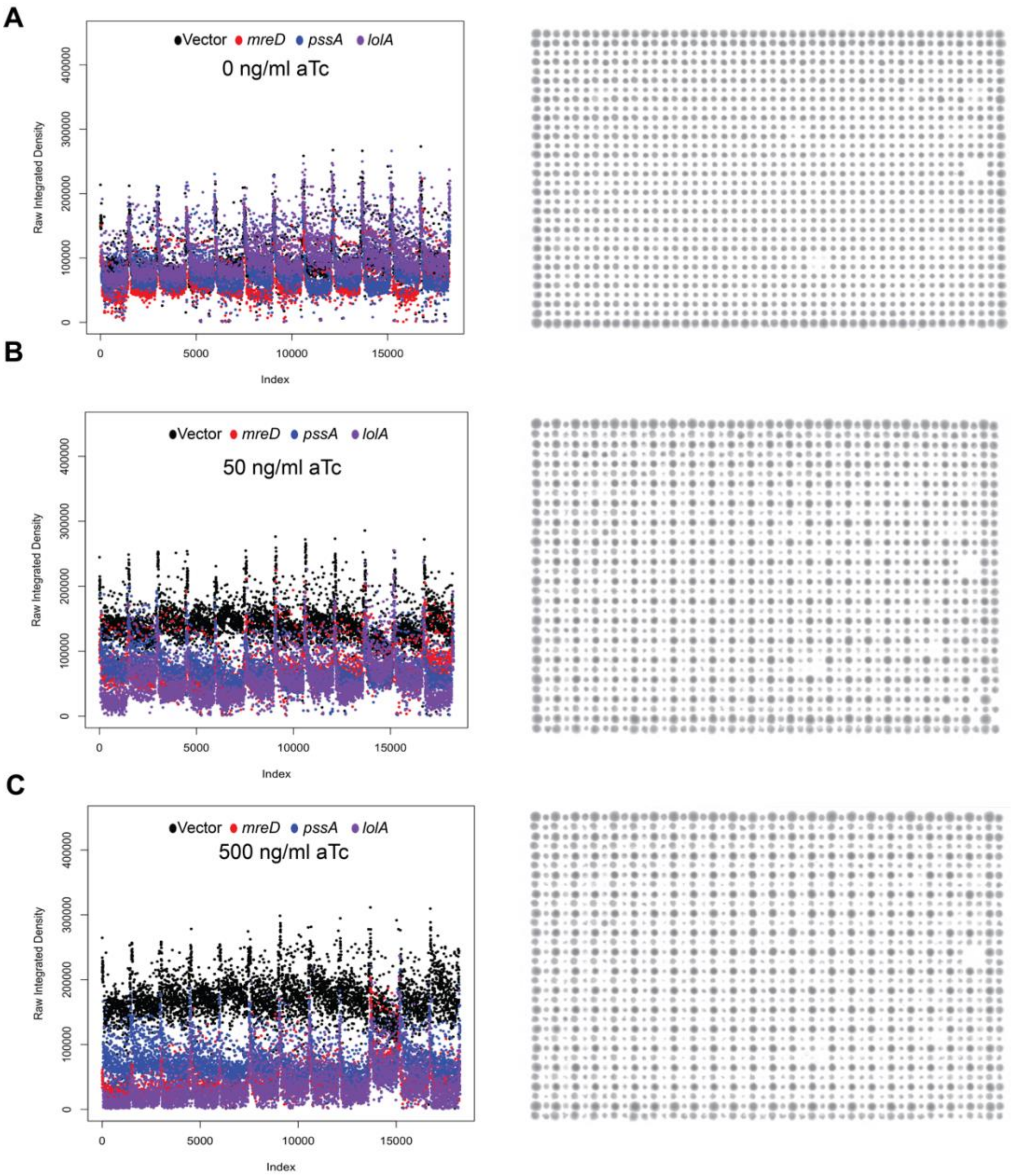


**Figure S8.** Representative index plots of raw integrated density and colony images of CRISPRi Keio cross at three different inducer concentrations: **(A)** 0 ng/ml, **(B)** 50 ng/ml and **(C)** 500 ng/ml of aTc. Shown is data for the cross plates containing the *mreD, pssA* and *lolA* targeting CRISPRi constructs. Colony images are shown from 384-well plate 4 of the Keio collection.

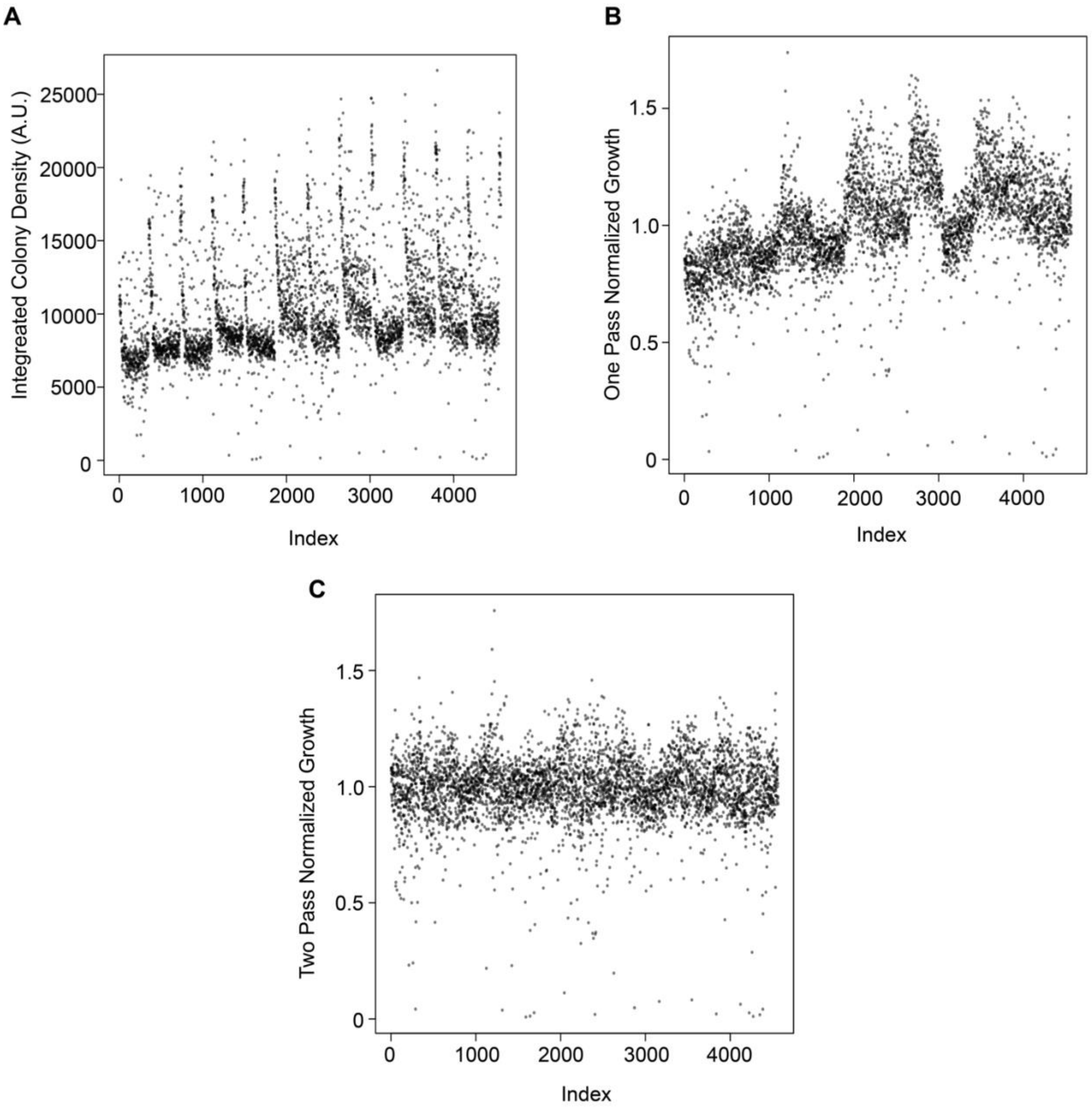


**Figure S9.** Normalization workflow for CRISPRi Keio screen, shown one replicate of the Keio pFD152_*lolA* dataset grown in the absence of aTc. Growth of the Keio collection is visualized by an index plot showing the growth of each strain in the collection. Shown here are plots showing the integrated density of each colony or raw colony growth **(A)**, the onepass normalized colony growth which accounts for spatial effects on growth **(B)**, and finally the two pass normalized dataset which normalized for plate to plate variability in addition to spatial effects.

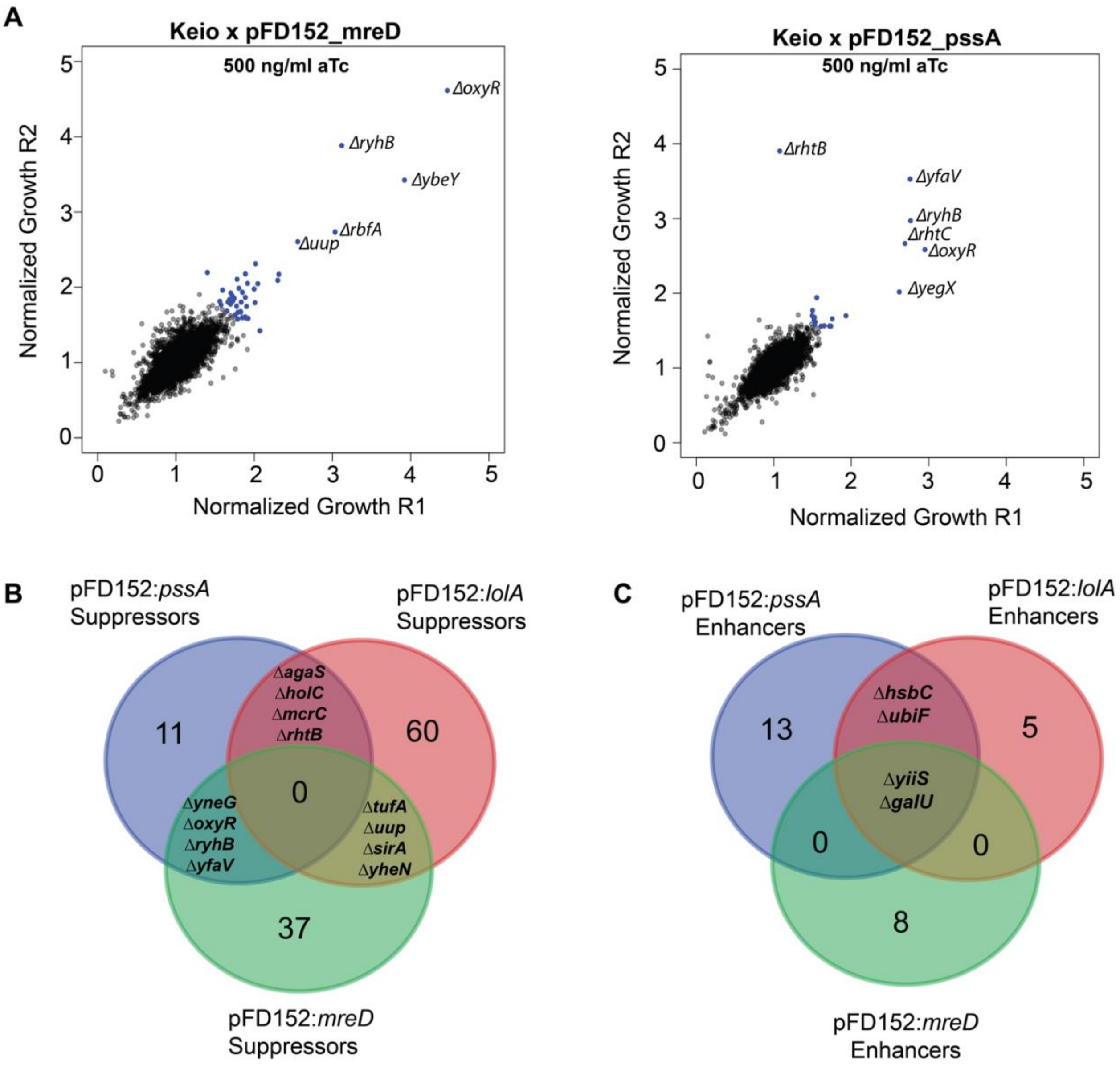


**Figure S10. (A)** Replicate plots of the *mreD* and *pssA* CRISPRi Keio screens at 500 ng/ml aTc. Mutants in blue demonstrate strains that differ from the mean of the dataset by at least three standard deviations. **(B)** Venn-diagram showing overlap of deletion mutants that suppress killing by CRISPRi at 500 ng/ml aTc from the three Keio crosses conducted. **(C)** Venn-diagram showing overlap of deletion mutants that enhance killing by CRISPRi constructs at 50 ng/ml from the three CRISPRi crosses conducted.
